# The Alfin-like transcription factor GmAlfin09 regulates endoplasmic reticulum stress in soybean via a peroxidase GmPRDX6

**DOI:** 10.1101/2023.02.05.527234

**Authors:** Kai Chen, Dongdong Guo, Jiji Yan, Huijuan Zhang, Zhang He, Chunxiao Wang, Wensi Tang, Yongbin Zhou, Jun Chen, Zhaoshi Xu, Youzhi Ma, Ming Chen

## Abstract

Soybean is a valuable oil crop cultivated throughout the world, but also highly susceptible to environmental stressors. The development of approaches to enhancing soybean stress resistance is thus vital to improving yields. In prior studies, Alfin has been shown to serve as an epigenetic regulator of plant growth and development. No studies of Alfin have yet been reported in soybean, however. In this study, the endoplasmic reticulum (ER) stress- and reactive oxygen species (ROS)-related transcription factor GmAlfin09 was identified. Screening of genes co-expressed with *GmAlfin09* unexpectedly led to the identification of the peroxidase GmPRDX6. Further analyses revealed that both *GmAlfin09* and *GmPRDX6* were responsive to ER stress, with GmPRDX6 localizing to the ER under stress. Promoter binding experiments confirmed the ability of GmAlfin09 to directly bind the *GmPRDX6* promoter. When *GmAlfin09* and *GmPRDX6* were overexpressed in soybean, enhanced ER stress resistance and decreased ROS levels were observed. Together, these findings suggest that GmAlfin09 can promote the upregulation of *GmPRDX6*, which subsequently localizes to the ER, reduced ROS levels, promotes ER homeostasis, and helps ensure the normal growth of soybean even under ER stress. This study highlights a novel genetic target for the future molecular breeding of stress-resistant soybean cultivars.

**HIGHLIGHT:** GmAlfin09 can increase the expression of *GmPRDX6* to reduce ROS level under ER stress.

## INTRODUCTION

*Alfin1*-like transcription factors were initially identified as salt stress-responsive genes in salt-tolerant alfalfa (Winicov I., 1993). The *Alfin1*-like gene family is highly conserved across species, with members being composed of 4 introns and 5 exons (Tuller et al., 2013). These Alfin1-like family proteins also exhibit a conserved N-terminal Alfin domain and a C-terminal PHD finger domain capable of specifically binding methylated H3K4me2/3 histones (Peña et al., 2006), thereby enabling these proteins to regulate chromatin accessibility and target gene transcription (Shi et al., 2006). These Alfin1-like proteins primarily exert their biological activities during the processes of seed germination, root growth and development, and root hair elongation (Chandrika et al., 2013; Goodrich et al., 2014). However, there is also clear evidence for the ability of Alfin1-like proteins to serve as important regulators of plant responses to abiotic and biotic stressors including exposure to cold, saline, drought, pathogen, and ABA stress (Wei et al., 2015; Swain et al., 2015; Janiak et al., 2016; Tao et al., 2018; Benny et al., 2019).

The endoplasmic reticulum (ER) is an organelle in which roughly one-third of all cellular proteins are synthesized. As such, ER-derived proteins play diverse roles in regulating the ability of cells, tissues, and organisms as a whole to respond to the surrounding environment (Reyes-Impellizzeri et al., 2021). Misfolded or unfolded proteins can begin to accumulate within the ER in response to a range of abiotic or biotic stressors, thereby subjecting cells to ER stress. Under physiological conditions, reactive oxygen species (ROS) function as signaling intermediaries and maintain redox equilibrium in addition to shaping stress responses. Recent work has shown that ROS production and redox metabolism are closely linked to ER stress such that insufficiently managed ER stress can enhance ROS production to a deleterious degree and subject plants to oxidative stress. However, ER stress can also activate nicotinamide adenine dinucleotide phosphate (NADPH) oxidase-mediated ROS signaling, bolster cellular antioxidant defenses, and alter the redox state of glutathione (GSH). ROS accumulation also plays a role in promoting ER stress response induction, suggesting a series of feedback mechanisms supported by evidence in *Arabidopsis* that ER stress can directly control ROS signaling, redox status, and antioxidant defense mechanisms (Ozgur et al., 2014). Plants have evolved a range of mechanisms to regulate ROS production and activity in response to ER stress conditions (Cao et al., 2022). When ER stress is not effectively reversed, however, this can irreversibly damage affected plants. The precise mechanisms responsible for the scavenging of ER stress-induced ROS, however, have yet to be fully clarified.

The bZIP, NAC, and WRKY transcription factors have been established as the primary regulators of ER stress responses. The majority of these proteins are ER-associated factors that must translocate to the nucleus to exert their biological functions. Members of the bZIP transmembrane family function by detecting ER stress, slicing itself C’ domain, transporting to nucleus and regulating the activation of proteins (Bailey and O’Hare, 2007). AtbZIP28 (Sun et al., 2015) or OsbZIP60 (Hayashi et al., 2013) is functionally analogous to mammalian ATF6 as the ER stress regulator. DTT and azetidine-2-carboxylate, which are drivers of the ER stress response, can also activate the AtbZIP60 (Iwata and Koizumi, 2005). The MaizeGDB database was used to identify the ER stress transducer AtbZIP17, which was designated ZmbZIP17 and found to be inducible in response to ABA treatment or exposure to ER stress-inducing agents including dithiothreitol (DTT) and tunicamycin (TM) (Yang et al., 2013). AtNTL7 is a membrane-tethered NAC transcription factor that is involved in resistance to ER stress in Arabidopsis (Chi et al., 2017). Under ER stress conditions, OsbZIP74 and OsNTL3 can also reportedly facilitate communication between the ER, plasma membrane, and nuclear compartments (Lu et al., 2012; Liu et al., 2020). The OsHLP1-OsNTL6 module is similarly involved in ER signal network formation and ER stress responses (Meng et al., 2022). ER stress-driven *OsWRKY45* expression depends on OsIRE1-OsbZIP50 pathway activity. AtWRKY33 can also be induced by ER stress (Hayashi et al., 2012). Established ER stress signaling models are likely to be incomplete, and the growing number of characterized transmembrane transcription factors associated with these responses highlight their selectivity and the complexity of their regulation (Bailey and O’Hare, 2007). No reports to date have demonstrated the ability of transcription factors to directly regulate ROS clearance in the context of ER stress.

Soybean (*Glycine max* (Linn.) Merr.) is an economically important crop species that serves as an important precursor for oil production and an important dietary source of protein. Here, a systematic analysis of soybean *Alfin* family genes was conducted. The roots are the first organ to respond to most abiotic stressors. Consistently, the expression levels of *GmAlfin9* were found to be highest in soybean roots, and to be responsive to the ER stress-inducing agent DTT. Functional analyses also demonstrated that GmAlfin9 can serve as a positive regulator of soybean ER stress responses. Co-expression and DNA binding analyses further revealed that GmAlfin9 was able to regulate the expression of *GmPRDX6*, which encodes a peroxidase capable of eliminating ROS in response to DTT treatment. In summary, these results demonstrate the ability of Alfin-like transcription factors to reduce the effects of abiotic stress on ER functionality in soybean cells through the upregulation of an ER-targeted peroxidase protein capable of eliminating ROS and restoring ER homeostasis. These results provide a more detailed understanding of the mechanisms used by soybean cells to alleviate ER stress while also offering novel targets for the future molecular breeding of stress-resistant soybean varieties.

## MATERIALS AND METHODS

### Plant materials, cultivation, and treatment

The W82 and mutant *gmalfin09* soybean cultivars were obtained from iSoybean (http://www.isoybean.org) (Zhang et al., 2022) established by Professor Qingxin Song of Nanjing Agricultural University. All materials were planted in a climate chamber at 28°C with a 16/8 h dark/light cycle.

ER stress was induced by treating plant cells with dithiothreitol (DTT) (Yang et al., 2013). Disulfide bond formation is a key step for the biosynthesis of many proteins, and the reducing agent DTT interferes with the oxidizing conditions necessary for these bonds to form in the context of protein folding, similarly causing misfolded proteins to accumulate in both the ER and cytosol. To focus on the role of disulfide bond formation, DTT (D104860, Aladdin, China) was used to treat plants in this study at a final 20 mM concentration by adding 20 mL of DTT treatment solution to the roots of plants while they were being cultivated in soil (Fig. 2A, Fig.6A). For plants subject to hydroponic cultivation, the DTT concentration in the nutrient solution was maintained at 20 mM (Fig. 6A).

**Figure 1.**
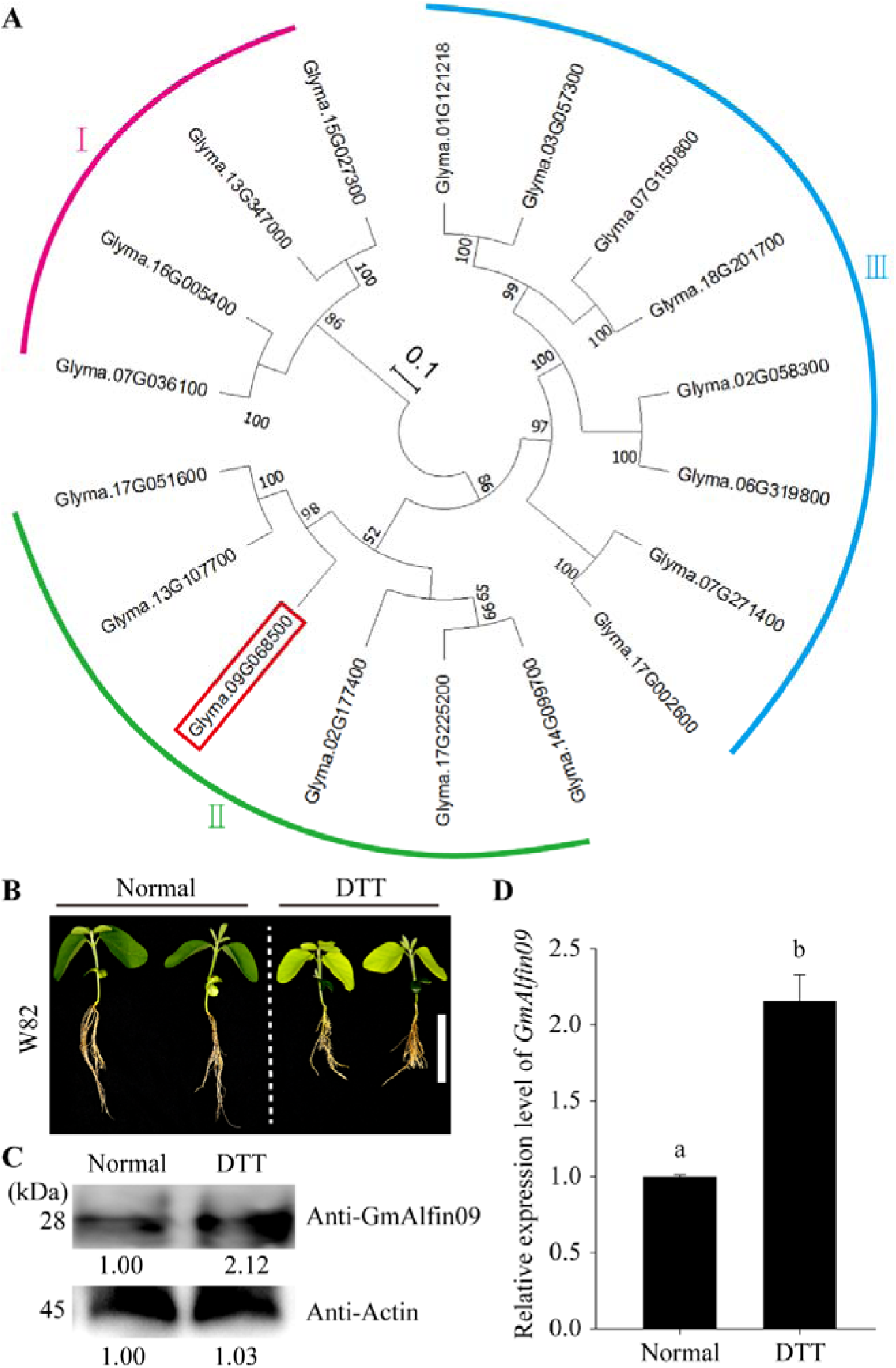
Phylogenetic tree construction and gene expression profile of *GmAlfin09*. **(A)** The construction of evolution tree of *GmAlfin* gene family. All protein sequences were shown in the Supplementary sequences. The evolutionary tree was displayed in a circle. Different subfamilies were represented by curves of different colors. Magenta represents the first sub family, green represents the second sub family, and blue represents the third sub family. The number at the resin node represents the bootstrap value. Scale bar represents sequence difference 0.1. The bootstrapped phylogenetic tree was constructed by using the MEGA-X software. The red box represents GmAlfin09 (Glyma.09G068500). **(B)** Phenotype of common cultivated soybean variety W82 (Seedling age is 3 days) under endoplasmic reticulum stress for 5 days simulated by 20mM DTT. The left side is the control group under normal growth conditions. Bar=10cm. **(C)** Analysis of protein GmAlfin09 expression level by western-blot using customized antibody. Take Actin protein as the sample uploading control. The gray value is below the strip. **(D)** The relative expression level of *GmAlfin09*. Statistically significant differences represented by different letters are derived from three biological repeats (ANOVA-DUNCAN’s multiple range tests at P<0.05). *GmCYP2* (Glyma.12g024700) gene was used as control. The error bars represent the SE of the mean. n=2×3.

**Figure 2.**
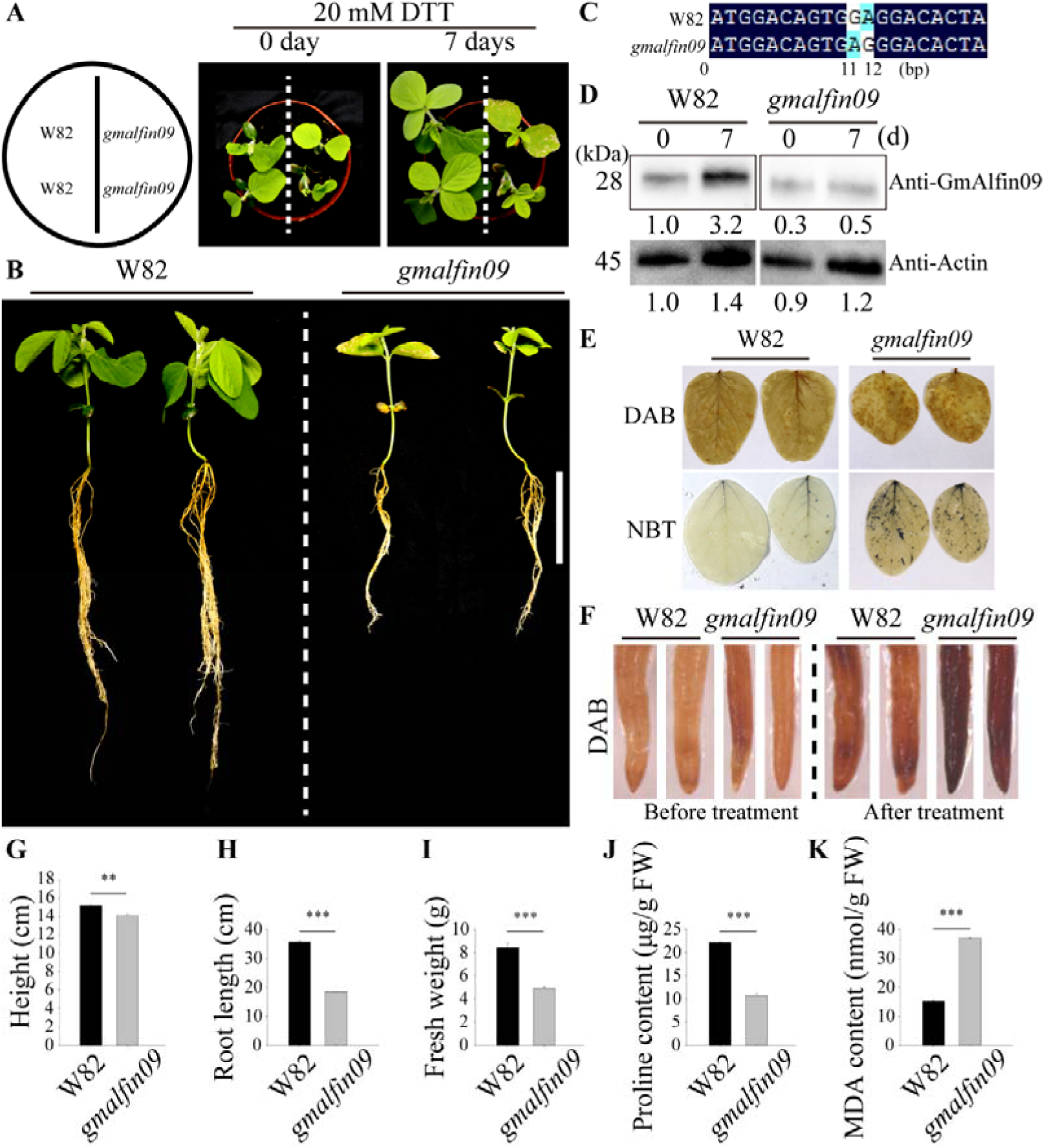
GmAlfin09 is necessary for soybean to resist ER stress. **(A)** This is a top view. Diagram (left) and observation (right, 0-7days) of ER stress treated cultivar W82 and mutant *gmalfin09* (Seedling age is 5 days) with 20mM DTT. **(B)** Display of the above ground and underground parts after treatment of Fig.1A. W82 is on the left, and *gmalfin09* is on the right. Bar, 10cm. **(C)** Comparison results of W82 and *gmalfin09* genomic sequences by software DNAMAN. The number below indicates the base position. **(D)** Before and after treatment, the protein content of GmAlfin09 in W82 (left) and *gmalfin09* (right) was displayed by western-blot using customized antibodies. The top is GmAlfin09, and the bottom is Actin. Take Actin protein as the sample uploading control. **(E)** After treatment, the leaves of W82 (left) and *gmalfin09* (right) were stained with DAB (above) and NBT (below). **(F)** DAB staining of W82 and *gmalfin09* root tips before and after treatment. **(G)** Plant height measured after treatment. **(H)** Root length measured after treatment. **(I)** Fresh weight measured after treatment. **(J)** Proline content measured after treatment. **(K)** MDA content measured after treatment. Data statistics conducted Student’s *t*-test, error bars represent SE of the mean, **P<0.01, ***P<0.001, n=4.

During phenotypic identification of mutant plants (Fig. 2A), soil-related interference was eliminated by planting all materials in the same pot and watering them regularly until two true leaves were present, at which time they were subject to DTT treatment by dripping 20 mL of a 20 M DTT solution on the base of the stem of each seedling until the soil had been fully saturated. The basic hydroponic culture medium used for this study was liquid 1/2MS_0_ culture medium (M519, Phyto Technology, USA), with DTT added to a final 20 mM concentration.

### Primers

Primers used for this study are listed in Supplementary Table S1.

### Immunoblotting

Samples of root and leaf tissue were initially homogenized in liquid nitrogen prior to the extraction of protein from these samples using an appropriate digestion buffer (50 mM Tris-HCl, pH=7.5, 150 mM NaCl, 0.1% NP-40, 4 M carbamide, 1 mM PMSF) (Qiao et al., 2012). Protein levels were then quantified using a Coomassie protein assay kit (B8522, Sigma-Aldrich, USA) followed by 12% SDS-PAGE separation (Bio-Rad, 1610175, USA) and immunoblotting with custom monoclonal mouse antibodies specific for GmAlfin09 (BPI, China) and GmPRDX6 (BPI, China) or with monoclonal mouse anti-Actin (plant specific) (ABclonal, AC009, China).

### Measurements of root length and plant weight

Roots were scanned using an Expression 11000XL root system scanning analyzer (Epson, Nagano, Japan) with a transparency adapter that had undergone calibration performed by Regent Instruments, Inc. Roots were arranged on the bed of the scanner so as to maximize secondary root separation while minimizing overlap. After scanning, root images were assessed to determine root length. Root data were analyzed with the WinRHIZO 2016 program (Regent Instruments, Inc., Quebec, Canada) (Kraft and Boge, 2001; Chen et al., 2022). To measure plant fresh weight, soil was washed off of the treated plants, after which excess water was removed with absorbent paper, and plants were weighed with a BSA224S-CW 1/10,000 analytical balance (Sartorius, Beijing, China). Three biological replicate samples were analyzed for these assays (Chen et al., 2022).

### DAB and NBT staining

Following exposure to control or DTT stress conditions, the roots and leaves of the treated seedlings were individually stained. For DAB staining, these samples were immersed for 12 or 24 h in DAB reagent (CSB-K09758AB-2; CUSABIO, Wuhan, China) and then transferred into 95% ethanol for decolorization. For NBT staining, samples were immersed in NBT stain (AR-0632; DINGGUO, Beijing, China) until dark blue (∼10 min for roots, ∼3 h for leaves) followed by decolorization with 95% ethanol. After staining, samples were imaged using a stereomicroscope (SZX16; OLYMPUS, Beijing, China) (Chen et al., 2022).

### Measurements of MDA and proline content

MDA and proline levels were analyzed in whole control (W82) or transgenic seedlings that had been exposed to DTT stress. Briefly, after seedlings had been subjected to DTT treatment for the appropriate number of days they were collected, and MDA levels were measured using an approach published previously (Du et al., 2018; Zhang et al., 2019; Chen et al., 2020). Briefly, ∼0.1 g plant tissue samples were collected for measurements of optical density (OD) at 532 and 600 nm. MDA levels were then calculated with the following formula: MDA content (nmol/g FW) = 51.6 × (OD532–OD600)/0.1. Proline levels were also determined using an approach published previously (http://www.cominbio.com/uploads/soft/180727/1-1PHGF414.pdf) after measuring the OD_520 nm_ values from four replicate ∼0.1 g samples of leaf tissue using the following formula: proline content (g/g FW) = 38.4 × (OD520 + 0.0021)/0.1. OD values were measured with a Varioskan LUX Multimode Microplate Reader (Thermo Fisher Scientific, USA). Analyses of three samples from the sample location were used when analyzing each transgenic line (Zhang et al., 2019; Chen et al., 2022).

### Transcriptional activation analyses

To confirm the transcriptional activation ability of GmAlfin09, the full-length *GmAlfin09* (GeneID: Glyma.09G068500) CDS sequence was linked to the pGBKT7 vector Gal4-DNA binding domain. The pGBKT7-*TaNAC48* (Chen et al., 2021) construct was selected as a positive control, and Y2Hgold yeast were transformed with these fusion vectors or an empty control vector followed by culture on SE/-Trp/-His/-Ade medium containing X-α-Gal (Chen et al., 2022).

### Subcellular localization assay

The full-length *GmAlfin09* and *GmPRDX6* (GeneID: Glyma.08G088000) CDS sequences without the stop codon were linked to the GFP sequence in the transient p16318hGFP expression vector, with the empty vector serving as a control. Soybean mesophyll cell protoplasts were transformed using TaNAC48-RFP and OsSD37-mCherry as nuclear and ER markers, respectively (Zhang et al., 2014). After they had been cultured in the dark for 20 h, a confocal laser microscope (ZEISS, LSM980 with Airyscan 2, Germany) was used to image these samples, and the ZEISS ZEN3.7 software was used to process a fluorescence density map for the resultant images.

### Measurements of chlorophyll content, H_2_O_2_ content, and conductivity

After treatment was complete, 4 representative leaves were selected from the same position of each plant, the veins were removed, and samples were cut into pieces, mixed, and 0.2 g was transferred into a 25 mL tube to which 10 mL of 80% ethanol was then added. The tube was then closed and transferred to a rotator for 24 h at room temperature until the leaves had faded. the treatment. Additional 80% ethanol was then added to a final volume of 25 mL, with the absorbance of the resultant chlorophyll extract solution being analyzed at 664 and 645 nm in a 1 cm diameter cuvette, with 80% ethanol serving as a blank control. Data were then analyzed as follows: C_T_=20.21A_645_+8.02A_663_, where C_T_ is the total chlorophyll concentration (mg/L); and Chlorophyll content (mg/g·FW) =C_T_×25(total extract volume)/0.2(sample fresh weight) ×1000.

An H_2_O_2_ Colorimetric Assay Kit (Beyotime, Shanghai, China) was used to quantify H_2_O_2_ levels in samples. REC testing was conducted as in a prior report published by Yu et al. (2006). Briefly, roots were washed and cut into small ∼1 cm pieces before addition to a tube containing 5 mL of deionized water. They were then incubated for 2 h at 25°C with occasional shaking, after which the electrical conductivity of the solution (C1) was measured. Samples were then boiled for 10 min, allowed to cool to room temperature (17–20°C), and conductivity was again measured (C2). REC was assessed with the following formula: REC (%) = C1/C2 9 100 (Li et al., 2008; Chen et al., 2021).

### qPCR

RNA was extracted from soybean plants in the experimental treatment groups using an RNA extraction kit (Plant RNA kit, TianGen, China), after which a Trans-Script One-Step RT-PCR SuperMix kit (Transgen, China) was used based on provided directions to prepare cDNA while removing genomic DNA. The TransStart Top Green qRT-PCR SuperMix (Transgen, China) was then used for qPCR analyses of gene expression in 96-well plates using the following settings: 45 cycles of 95°C for 15 s, 60°C for 20 s, and 72°C for 45 s (Dong et al., 2016). Primers used for this assay are listed in Supplementary Table S1. Soybean actin served as a normalization control gene. Data are reported as means ± SE from three replicate samples (Chen et al., 2022).

### *Agrobacterium rhizogenes* transformation of soybean hairy roots

The *GmAlfin09* and *GmPRDX6* genes amplified and cloned into the pCAMBIA1302, with subsequent transformation then being performed as reported by Kereszt et al., (2007).

### GUS staining

Following the amplification of the *GmAlfin09* and *GmPRDX6* promoter fragments (−2 kb from the ATG site), they were cloned into the pCAMBIA1305 vector, after which soybean hairy root transformation was performed as reported by Kereszt et al. (2007). Tobacco leaves were transiently transformed as reported previously (Yu et al., 2021), using recombinant vectors that were transformed into competent *A. rhizogenes* GV3101 cells (Zoman Biotechnology). *A. rhizogenes*-mediated transformation was then used to co-express recombinant and blank control vectors in *N. benthamiana* leaves for 3 days, after which these leaves were removed and subjected to surface treatment with DTT for 12 h while protected from light. Samples were then stained for 3 h in GUS staining buffer (50 mM sodium phosphate [pH 7.0], 10 mM EDTA-Na_2_, 20% methanol, 0.1% Triton X-100, and X-Gluc at 0.5 mg/ml) at 37°C following vacuum infiltration for 5 min. Experiments were repeated in triplicate, and greater than 10 *GmAlfin09 pro:*:GUS or *GmPRDX6 pro:*:GUS transgenic plants were imaged with a ZEISS Stereo Discovery.V20 instrument (Jena, Germany).

### Electrophoretic mobility shift assay

A LightShift® Chemiluminescent EMSA Kit (Thermo Fisher Scientific) was used based on provided directions (Tang et al., 2021). Briefly, *GmAlfin09* coding regions were cloned into the GST tag-containing pGEX4T-1 vector, with recombinant GST-GmAlfin09 then being induced using 1 mM IPTG followed by expression in the *Escherichia coli* Rosetta (DE3) strain. Glutathione-Sepharose^TM^ 4B beads (GE Healthcare, IL, USA) were used for recombinant protein purification, and these proteins were then quantified using a BCA Protein Assay Kit (Sangon Biotech, C503021). Oligonucleotide probes were subject to 5’ biotinylation, while competitor sequences for binding assays were unlabeled (Supplementary Table S1). Equal amounts (1µl) of GST-GmAlfin09 fusion proteins were incubated for 20 min in a total volume of 20 µL of binding buffer (50 ng·ml/L Poly(dI-dC), 2.5% glycerol, 0.05% NP-40, 10 mM EDTA, 0.5 mM DTT, and 20 pmol probes) at room temperature. These solutions were then loaded onto 6% polyacrylamide gels and separated at 4°C with 0.5× Tris-Borate-EDTA buffer at 4°C. DNA–protein complexes were then transferred to a nylon membrane (Thermo Fisher Scientific). UV cross-linking was performed, after which biotin was detected based on provided directions (Thermo Fisher Scientific) (Tang et al., 2021).

### Dual-luciferase activity assays

Dual-LUC activity assays were conducted using soybean mesophyll cell protoplasts and the pGreen II 0800-LUC vector system that includes the firefly luciferase (LUC) gene under the control of the promoter sequence for the target gene of interest and the control Renilla luciferase (REN) gene under the control of the constitutive 35S promoter (Tang et al., 2021). The HindIII and BamHI sites were used to clone the amplified *GmPRDX6* promoter fragment (−2 kb from the ATG site) into the pGreenII 0800-LUC vector to generate the *GmPRDX6 pro::LUC* vectors. The targeted mutation of this promoter sequence was achieved using two overlapping primer pairs with mutation sites containing the mutated cis-elements (CEs), respectively generating *mGmPRDX6 pro^p1^::LUC*, *mGmPRDX6 pro^p2^::LUC* and *mGmPRDX6 pro^p1p2^::LUC* vectors (Tang et al., 2021). The p16318hGFP (GmAlfin09-GFP) construct served as an effector. Transformation was performed as in subcellular localization assays. LUC activity for plant extracts was assessed with a microplate reader and commercial LUC reaction reagents based on provided directions (Promega, WI, USA) (Tang et al., 2021).

## RESULTS

### *GmAlfin09* is an ER stress-inducible member of *GmAlfin* gene subfamily II

As there have been no prior reports of the soybean *Alfin* gene family, to comprehensively survey these genes an initial analysis of *GmAlfin* genes throughout the soybean genome was conducted (Fig. 1A). This approach identified 18 *GmAlfin* genes, the corresponding protein sequences (Supplementary sequences) for which were then introduced into the Mega-X software to generate a Neighbor-Joining phylogenetic tree. This approach clustered these 18 proteins into three main subfamilies (I, II, and III). As the subfamily II member Glyma.09G068500 was encoded on soybean chromosome 9, it was tentatively named *GmAlfin09*.

To more fully characterize the *GmAlfin* gene family, expression data for this gene family (Supplementary Table S2) were analyzed with HemI 1.0 (Supplementary Fig. S1A). This approach revealed that with the exception of a limited number of these genes (including Glyma.02G058300, Glyma.02G177400, and Glyma.06G319800), the majority of genes in the *GmAlfin* family were expressed at higher levels in root and seed tissues, with *GmAlfin09* expression levels being highest in the roots, root tips, and nodules (Supplementary Fig. S1A). *GmAlfin09* expression was further visualized in a range of tissues and organs using Plant eFP (http://bar.utoronto.ca/eplant_soybean/) (Supplementary Fig. S1B), and a phenotypic analysis of the common W82 soybean cultivar was conducted (Fig. 1B). This assay revealed that DTT treatment suppressed soybean plant growth with concomitant root length shortening (Fig. 1B). GmAlfin09 protein levels rose in response to DTT treatment as detected with a custom monoclonal mouse antibody (Fig. 1C), consistent with the observed increase in *GmAlfin09* levels (Fig. 1D). Given the importance of roots for the detection of and resistance to external stressors, these high root expression levels of *GmAlfin09* were of particular interest such that this gene was selected as the target for all subsequent research in this manuscript.

### *GmAlfin09* is required for ER stress resistance in soybean

In a prior study, we obtained a mutated *GmAlfin09* sequence (hereafter *gmalfin09*) from the iSoybean database used for the mutational fingerprinting of soybean plants (Zhang et al., 2022). Phenotypic analyses revealed that *gmalfin09* was associated with a significant reduction in ER stress resistance (Fig. 2). To ensure that this result was not attributable to heterogeneous soil conditions, control W82 and *gmalfin09* plants were simultaneously planted in the same pot with two biological replicates (Fig. 2A). Prior to any treatment, *gmalfin09* grew more slowly than wild-type W82 plants. When two true leaves were present on these seedlings, treatment was then initiated after which time the leaves of *gmalfin09* plants became withered, shrunken, and yellow on day 7 following DTT (20 mM) treatment (Fig. 2A). After washing away root-associated soil, *gmalfin09* root development was also visibly weaker than that of W82 seedlings (Fig. 2B). Significant differences in W82 and *gmalfin09* plant height were also observed (Fig. 2G), with extremely significant differences in root length and fresh weight (Fig. 2H-I). Sequence alignment identified two bases in the CDS region that differed between the W82 and *gmalfin09* sequences together with the substitution of a single amino acid at the protein level (Fig. 2C; Supplementary Fig. S1C). This amino acid substitution was not predicted to alter the overall protein conformation or the conformation of the main functional PHD domain (Supplementary Fig. S1C). Even so, GmAlfin09 protein levels were clearly affected by this substitution (Fig. 2D). A custom mouse monoclonal antibody was therefore used to detect changes in GmAlfin09 protein levels prior to and after treatment, revealing a lower basal GmAlfin09 level in the roots of *gmalfin09* plants relative to wild-type W82 plants, and these levels failed to rise in response to DTT. In contrast, GmAlfin09 levels in wild-type plants rose significantly upon treatment (Fig. 2D). Post-treatment proline levels in *gmalfin09* plants were significantly lower than those in W82 plants (Fig. 2J), whereas malondialdehyde (MDA) levels were significantly higher than those in wild-type plants (Fig. 2K). These results highly the close relationship between GmAlfin09 and ER stress.

### GmAlfin09 responds to ER stress by directly promoting *GmPRDX6* upregulation

The above results suggest that appropriate *GmAlfin09* expression is required for soybean plants to effectively respond to ER stress, but the underlying molecular mechanisms have yet to be established. To test the potential of GmAlfin09 to serve as a transcriptional activator, Y2HGold yeast transformation was next performed (Fig. 3A). This assay revealed that yeast strains transformed with *GmAlfin09* and the positive control *TaNAC48* were able to grow on SE/- Trp/- His/- Ade medium and show blue color (Fig. 3B), whereas those transformed with the lam control were unable to grow, confirming that GmAlfin09 exhibits transcriptional activation activity. To directly explore ER stress-induced changes in the localization of GmAlfin09, soybean protoplasts were transiently transformed with the *GmAlfin09-GFP* expression construct (Fig. 3C). Under basal conditions, GmAlfin09 was distributed in and around the nucleus (Fig. 3C-D). In response to DTT treatment, GmAlfin09 fully localized to the nucleus (Fig. 3C-D), suggesting that it can response to ER stress in nucleus whereupon it serves as a transcriptional activator. GUS staining was further performed to verify *GmAlfin09* responses to DTT (Fig. 3E). As *GmAlfin09* was expressed in the roots of soybean plants (Supplementary Fig. S1A-B), this assay was performed using the soybean hairy root transformation system (Kereszt et al., 2007). Deeper GUS staining was evident for *GmAlfin09 pro::GUS* samples, with higher levels of *GUS* gene expression in response to treatment with DTT (Fig. 3E). Quantitative analyses also confirmed the upregulation of the *GUS* gene in response to DTT induction (Fig. 3F). These data support the ability of *GmAlfin09* to respond to DTT-induced ER stress.

**Figure 3.**
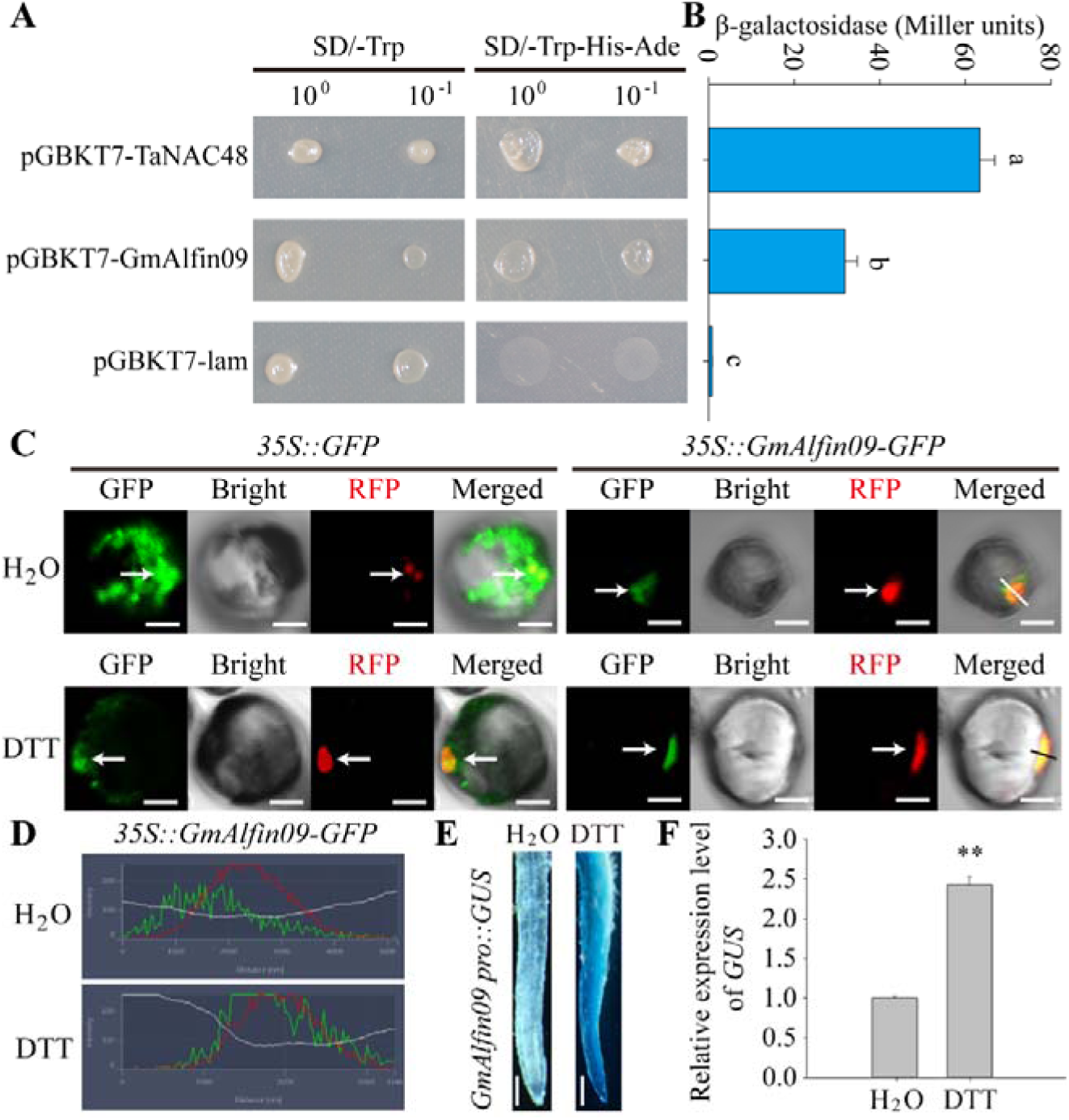
GmAlfin09 was identified as a transcription factor in response to ER stress. **(A)** Verification of activation ability of transcription factor GmAlfin09 by using yeast strain Y2HGold. pGBKT7-TaNAC48 was the positive control and pGBKT7-lam was the negative control. The result was judged by the growth of the colony on the plate medium SE/-Trp-His-Ade. The concentration on the left side was normal, and the concentration on the right side was diluted 10 times. The concentration on the left side is normal, and the concentration on the right side was diluted 10 times. **(B)** β-galactosidase activity in the cultures of yeast cells incubated with X-α-gal. Statistically significant differences represented by different letters are derived from three biological repeats (ANOVA-DUNCAN’s multiple range tests at P<0.05). The error bars represent the SE of the mean. n=3. **(C)** Subcellular localization of GmAlfin09 in soybean mesophyll protoplasts. The left side was the empty control (*35S::GFP*), and the right side was the localization of GmAlfin09. The lower row was the result of 20mM DTT treatment, and the upper row was replaced by water. RFP was nuclear marker. The white arrow pointed to the nucleus. Bar=10nm. Merged views of GmAlfin09 were sampled, and the sampling ranges were indicated by a line. **(D)** Fluorescence density diagram within the sampling ranges in Fig. 3C measured by ZEISS lite. The green line represents GFP fluorescence, and the red line represents the red fluorescence of nuclear marker. **(E)** Promoter activity analysis in soybean hairy roots by GUS staining. The right side was the DTT treatment group, and the left side was replaced by water. Bar=0.1mm. After decolorization, the leaves were observed with ZEISS Stemi 508. **(F)** Relative expression level of *GUS* gene in soybean hairy roots by qRT-PCR. *GmCYP2* (Glyma.12g024700) gene was used as control. Data statistics conducted Student’s *t*-test, error bars represent SE of the mean, **P<0.001, n=3.

Next, the Phytozome database was used for *GmAlfin09* co-expression network analyses. This approach revealed strong correlations between *GmAlfin09* expression and the expression of several peroxidase genes (https://phytozome-next.jgi.doe.gov/report/gene/Gmax_Wm82_a4_v1/Glyma.09G068 500). In line with this observation, *gmalfin09* mutant plants were found to exhibit higher ROS levels (Fig. 2E-F). The DAB and NBT staining of leaf and root samples indicated that significantly higher H_2_O_2_ and superoxide anion levels were evident in *gmalfin09* leaves following treatment relative to W82 leaves (Fig 2E), with a similar increase in H_2_O_2_ levels in *gmalfin09* roots relative to W82 roots (Fig. 2F). To gauge the ability of these plants to resist oxidative stress, the proline and malondialdehyde levels in these plants were further analyzed (Fig. 2J-K), providing support for a model in which *GmAlfin09* regulates ER stress responses through the ROS signaling pathway.

Among the analyzed redox-related proteins, *GmPRDX6* was most strongly correlated with *GmAlfin09* in co-expression analyses (https://phytozome-next.jgi.doe.gov/report/gene/Gmax_Wm82_a4_v1/Glyma.09G068 500), suggesting that it may be associated with ER stress resistance. Analyses of the *GmPRDX* gene family indicated that *GmPRDX6* is a member of subfamily in this gene family (Supplementary Fig. S3A), with different protein structural characteristics in each of these three protein subfamilies. Tissue-level analyses of gene expression revealed the expression of *GmPRDX6* in soybean roots, leaves, and pods (Supplementary Fig. S3B). Roots of DTT-treated W82 plants were next used for quantitative analyses of the expression of all members of the *GmPRDX* gene family (Supplementary Fig. S3C). Promoter binding element analyses revealed that many Alfin binding sites were present in *GmPRDX* promoter regions (Supplementary Fig. S4), and qPCR analyses indicated that the relative expression levels of *GmPRDX6* were higher than those of all other members of this gene family in response to treatment (Supplementary Fig. S3C). These data indicated that *GmPRDX6* is critical for soybean ER stress responses and may be an important GmAlfin09 target gene.

To verify the ability of GmAlfin09 to serve as a direct regulator of *GmPRDX6*, EMSA and dual-luciferase reporter assay experiments were next performed. An initial analysis of the *GmPRDX6* promoter sequence identified two potential GmAlfin09 binding elements (Fig. 4A). In an EMSA assay, blocking bands were evident in the lanes incubated with GST-GmAlfin09, P1-biotin (Fig. 4B), or P2-biotin (Fig. 4C), but were weakened or absence following competitive probe addition or probe mutation (Fig. 4B-C). This suggests that GmAlfin09 can directly interact with the two identified binding sites in the *GmPRDX6* promoter region. Consistently, following the construction of plasmids necessary for a dual-LUC assay (Fig. 4D), GmAlfin09 was confirmed to bind to both the P1 and P2 promoter sites (Fig. 4D-F), exhibiting particularly strong binding to the P2 site (Fig. 4F).

**Figure 4.**
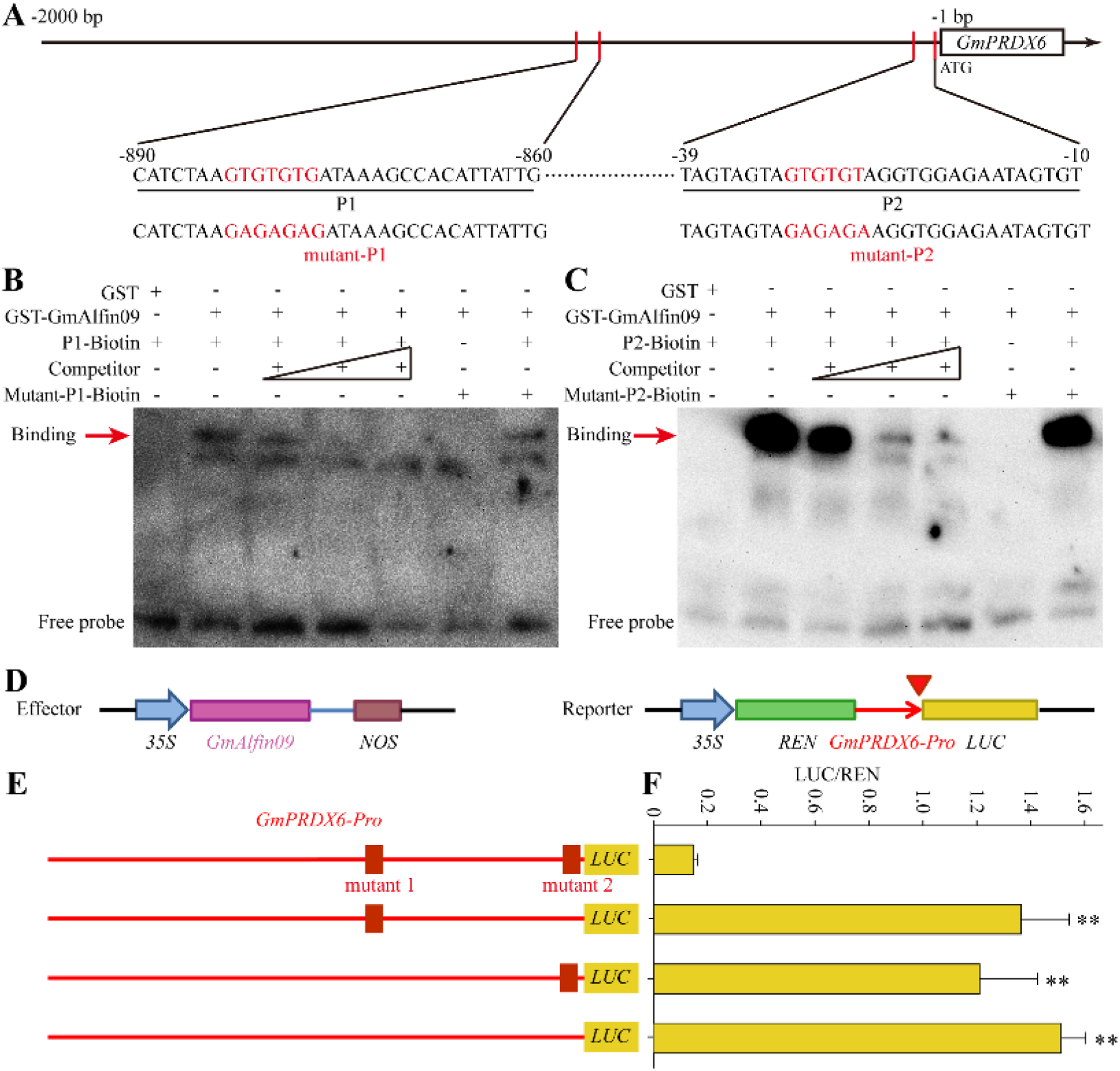
*GmPRDX6* is the target gene of GmAlfin09 direct binding. **(A)** Schematic structures of *GmPRDX6* promoter showing the presence of two binding sites [one site at -890∼-860bp (P1 site) and the other at -39∼-10bp (P2 site) from the ATG translation start codon] and labelled DNA probes in the Electrophoretic Mobility Shift Assay (EMSA). **(B-C)** EMSA analysis of protein-DNA complexes performed with recombinant GST-GmAlifin09 protein and a fragment of the promoter containing each of the two sites as a probe. Lane 1, GST/free probe; lanes 2, GST-GmAlifin09/free probe; lanes 3-5, protein-DNA complexes were competed out with a 10-fold molar excess of unlabelled oligonucleotide containing a consensus site (Competitor); lanes 6, GST-GmAlifin09/ free probe with mutanted oligonucleotide; lanes 7, specific GST-GmAlifin09/free probe protein-DNA binding was challenged with mutanted-probe. The red arrow represents the banding. **(D)** Schematic diagram of plasmid construction in dual-luciferase reporter assay (LUC). **(E)** Schematic diagram of the effect of GmAlfin09 on the expression of *GmPRDX6 pro*::*LUC* when transiently expressed in soybean protoplasts. (F) Data statistical analysis of LUC/REN values during the experiment. Asterisks indicate significant differences with respect to means of the control (Student’s *t*-test): **P < 0.01, n=3.

While GmAlfin09 can directly bind to the *GmPRDX6* promoter, further research is necessary to definitively show that *GmPRDX6* can participate in ER stress responses. Accordingly, GUS staining was performed to confirm the direct responsivity of *GmPRDX6* to ER stress based on an analysis of *GmPRDX6* promoter activity (Fig. 5). This approach confirmed that *GmPRDX6* responded to the DTT treatment of tobacco leaves (Fig. 5A-E). This analysis was also conducted in soybean hairy roots to more closely mimic physiologically relevant conditions, again demonstrating that *GmPRDX6* was responsive to the DTT treatment of soybean plants (Fig. 5F-G). The subcellular localization of GmPRDX6 under ER stress conditions was also visualized by transiently transforming soybean protoplasts with a *GmPRDX6-GFP* expression construct (Fig. 5H-I). This analysis revealed that GmPRDX6 was largely distributed in the cytosol at baseline (Fig. 5H-I). However, following DTT treatment, the GmPRDX6 fluorescent signal primarily overlapped with that of the ER (Fig. 5H-I). This thus suggests that GmPRDX6 can enter the ER to function as an oxidoreductase in response to ER stress.

**Figure 5.**
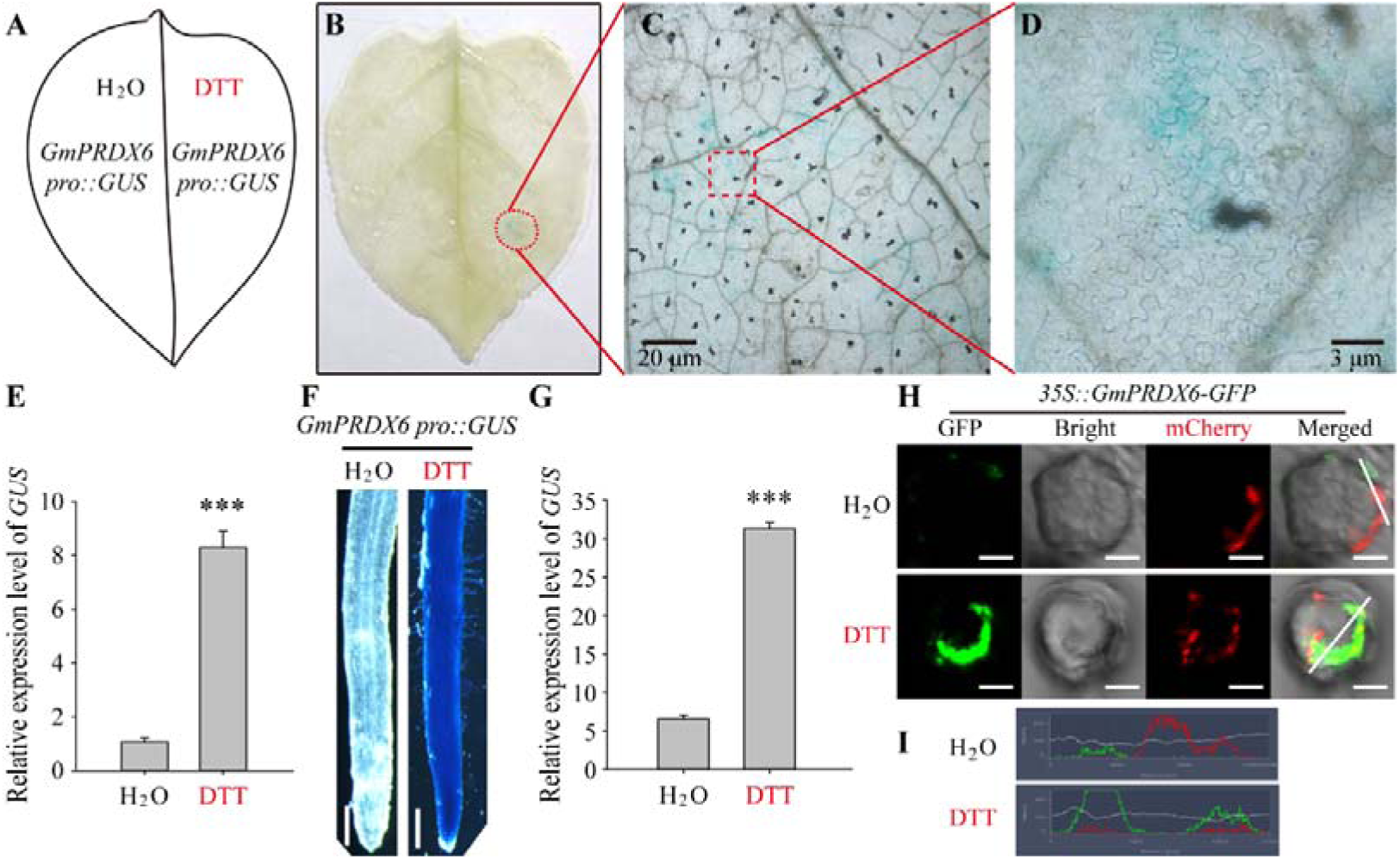
*GmPRDX6* can respond to the ER stress simulated by DTT. **(A-D)** Promoter activity analysis in tobacco leaves by GUS staining. **(A)** At the far left was the treatment diagram. The right side was the DTT treatment group, and the left side was replaced by water. **(B-D)** After decolorization, the leaves were observed with ZEISS Stemi 508. On the right were locally magnified views. **(E)** Relative expression level of *GUS* gene in tobacco leaves by qRT-PCR. *GmCYP2* (Glyma.12g024700) gene was used as control. Data statistics conducted Student’s *t*-test, error bars represent SE of the mean, ***P<0.001, n=3. **(F)** Promoter activity analysis in soybean hairy roots by GUS staining. The right side was the DTT treatment group, and the left side was replaced by water. Bar=0.1mm. **(G)** Relative expression level of *GUS* gene in soybean hairy roots by qRT-PCR. Data statistics conducted Student’s *t*-test, error bars represent SE of the mean, ***P<0.0001, n=3. **(H)** Changes of subcellular localization of GmPRDX6 in soybean protoplasts after DTT treatment. The lower row was the result of 20mM DTT treatment, and the upper row was replaced by water. mCherry was ER marker. Bar=10nm. Merged views of GmPRDX6 were sampled, and the sampling ranges were indicated by a line. **(I)** Fluorescence density diagram within the sampling ranges in Fig. 5H measured by ZEISS lite. The green line represents GFP fluorescence, and the red line represents the red fluorescence of ER marker.

### Verification of the function of the soybean GmPRDX6 protein

To validate the functions of the GmPRDX6 protein, *GmPRDX6-RNAi* knockdown and *GmPRDX6* overexpression lines were established via a soybean hairy root transformation approach (Fig. 6). Prior to treatment, the *GmPRDX6-RNAi* lines exhibited the poorest growth of the analyzed lines (Fig. 6A). During treatment, yellowing of the leaves and roots was observed for each of these lines. On day 10 of treatment, leaves for all lines began to wilt and the plants were returned to normal growth medium. Following a 5-day recovery period, the roots and leaves of the GmPRDX6 knockdown lines remained yellow and withered, whereas the stem tip leaves of plants overexpressing *GmPRDX6* overexpression lines began to turn green and the roots began to turn white (Fig. 6A). DAB and NBT staining results demonstrated that the deepest staining was evident in the leaves and roots of the *GmPRDX6-RNAi* knockdown lines consistent with the highest levels of H_2_O_2_ and superoxide anions therein (Fig. 6B-C). In contrast, the leaves and roots of plants overexpressing *GmPRDX6* exhibited the lightest DAB and NBT staining (Fig. 6B-C). qPCR analyses indicated that following treatment, relative *GmPRDX6* expression levels remained highest and lowest in the *GmPRDX6* overexpression and *GmPRDX6-RNAi* knockdown lines, respectively (Fig. 6D). Physiological analyses further confirmed that the highest root length, fresh weight, chlorophyll, and proline content values were observed in the *GmPRDX6* overexpression lines, whereas the MDA levels in these plants were lowest (Fig. 6E-I). These results suggest that overexpressing *GmPRDX6* can enhance the ability of soybean plants to resist DTT-induced ER stress.

**Figure 6.**
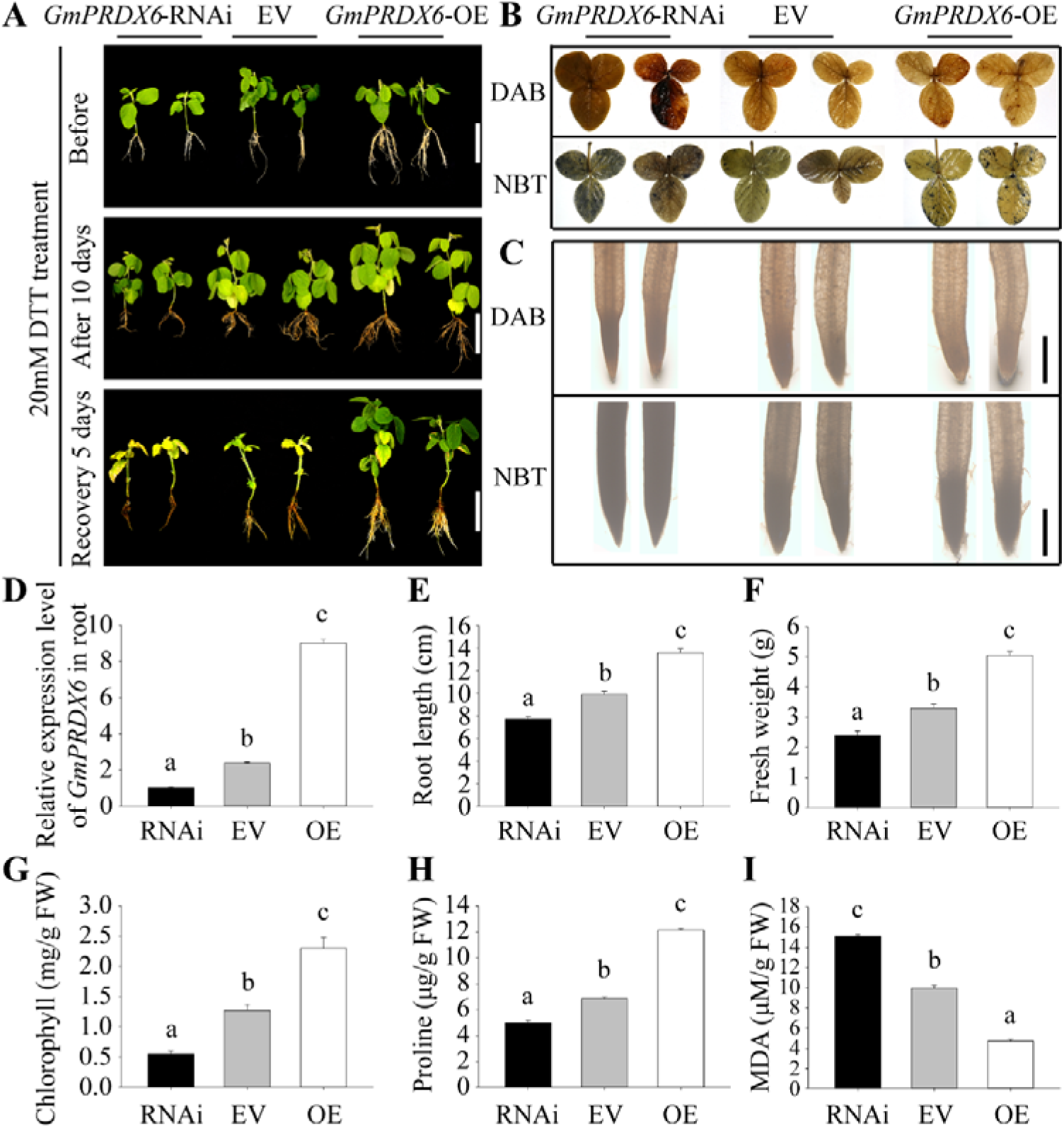
The expression of *GmPRDX6* gene is necessary for soybean to resist ER stress. **(A)** Phenotypes of plants with transgenic soybean hairy roots under 20mM DTT treatment. From left to right, the picture showed the RNAi interference lines of *GmPRDX6*, the empty vector control group (EV) and the overexpression lines of *GmPRDX6*. Bar=10cm. **(B)** DAB and NBT staining of leaves of each line after treatment. **(C)** DAB and NBT staining of hairy roots of each line after treatment. Bar=0.1mm. **(D)** Relative expression level of *GmPRDX6* in soybean hairy roots of each line. *GmCYP2* (Glyma.12g024700) gene was used as control. **(E)** Root length of each line. **(F)** Fresh weight of each line. **(G)** Chlorophyll content of each line. **(H)** Proline content of each line. **(I)** MDA content of each line. FW, fresh weight. **(D-I)** Data statistics conducted ANOVA-DUNCAN’s multiple range tests at P<0.05, error bars represent SE of the mean, n=3.

### Combined analysis of the functional roles of *GmAlfin09* and *GmPRDX6* during ER stress

To more directly assess the functional relationship between *GmAlfin09* and *GmPRDX6* in soybean plants, overexpression lines were established by transferring *GmAlfin09* or *GmPRDX6* into soybean hairy roots and then subjecting these transgenic plants to DTT treatment. While control plants transformed with an empty vector began to wild on day 3 of treatment, the *GmAlfin09*-OE line began to wilt on day 4, and the *GmPRDX6*-OE line began to wilt on day 5 (Fig. 7A). The root system of the *GmPRDX6*-OE line was the most developed, whereas that of the control group was the least developed (Fig. 7B). DAB and NBT staining of the leaves and root tips of these three soybean lines were next used to assess the ROS levels in each strain, demonstrating that this staining was lightest for the leaves and root tips of the *GmPRDX6*-OE line, followed by the *GmAlfin09*-OE line, with control plants exhibiting the deepest staining (Fig. 7C-D; Fig. 8). The fresh weight and root length of *GmPRDX6*-OE line were the highest among the three lines (Fig. 7E-F), whereas the H_2_O_2_ levels and conductivity levels were lowest (Fig. 7G-H). The physiological results for the *GmAlfin09*-OE line were between those for the *GmPRDX6*-OE line and the control plants (Fig. 7E-H). When Western immunoblotting was used to detect GmAlfin09 and GmPRDX6 protein levels in these plants, DTT treatment was confirmed to promote the upregulation of both of these proteins (Fig. 7I). Prior to treatment, GmAlfin09 protein levels were generally higher than those of GmPRDX6. However, the change in GmPRDX6 amplitude was greater than that for GmAlfin09. The GmAlfin09 protein level in the *GmAlfin09*-OE line rose the most quickly and to the greatest extent among the tested lines, while the same was true for GmPRDX6 in the *GmPRDX6*-OE line. In contrast, control plants exhibited lower levels of GmAlfin09 and GmPRDX6 expression (Fig. 7I).

**Figure 7.**
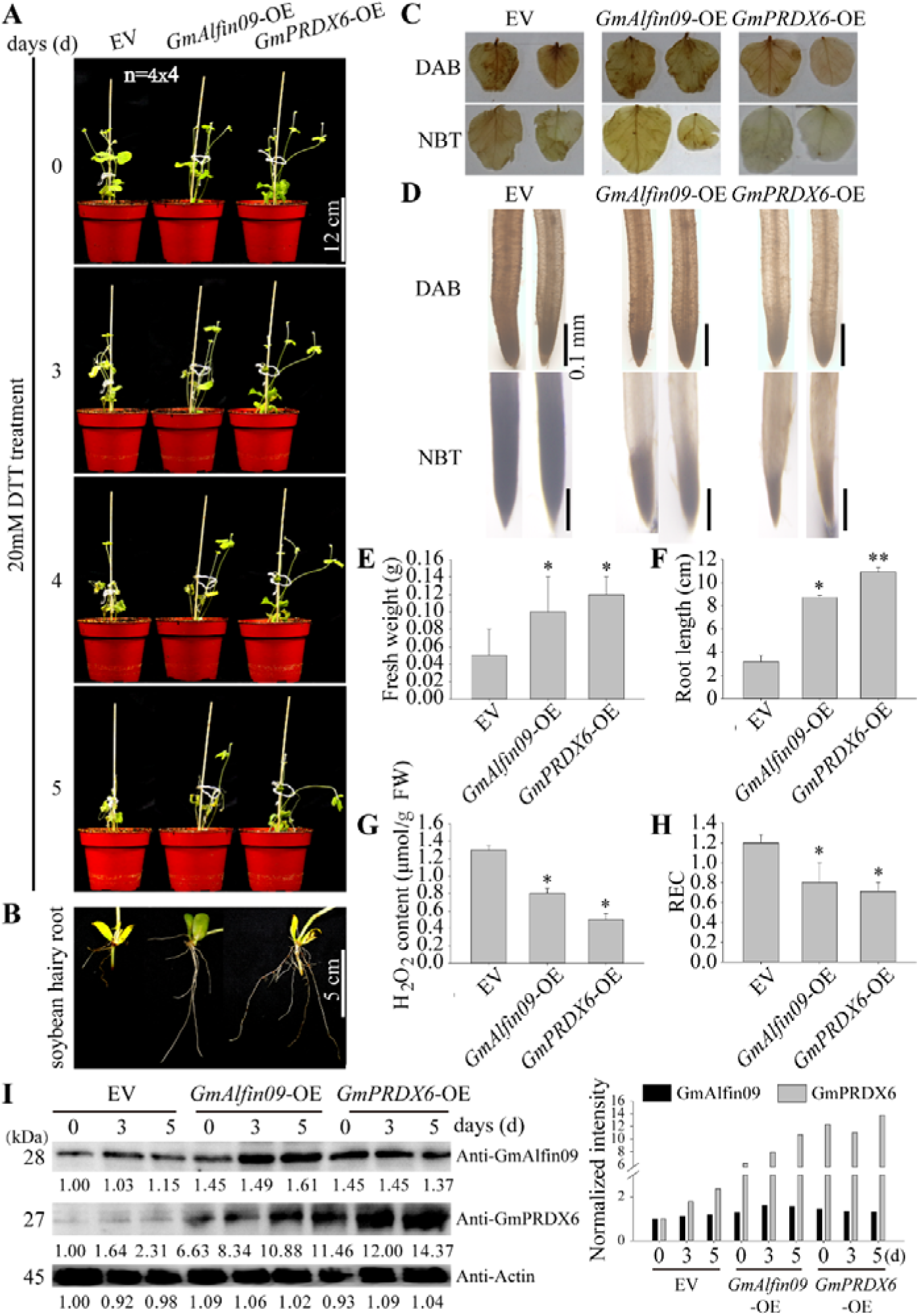
Continuous observation of phenotypes of *GmAlfin09* and *GmPRDX6* overexpression lines under ER stress. **(A)** The materials from left to right in the figure are the empty-vector control group (EV), the overexpression lines of *GmAlfin09* and *GmPRDX6*. Continuous observation was carried out, Bar=12cm. **(B)** The last state of the roots. Bar=5cm. **(C)** DAB and NBT staining of leaves of each line after treatment. **(D)** DAB and NBT staining of hairy roots of each line after treatment. Bar=0.1mm. **(E)** Fresh weight of each line. **(F)** Root length of each line. **(G)** H_2_O_2_ content of each line. **(H)** REC of each line. Data statistics conducted Student’s *t*-test, error bars represent SE of the mean, *P<0.01, **P<0.001, n=4 4. **(I)** The changes of protein level of each line during treatment. Customized antibody Anti-GmAlfin09 and Anti-GmPRDX6 were used. Take Actin protein as the sample uploading control. On the left is the western blot result, below the stripe is the grayscale value and on the far right is the histogram of the gray value of each stripe.

**Figure 8.**
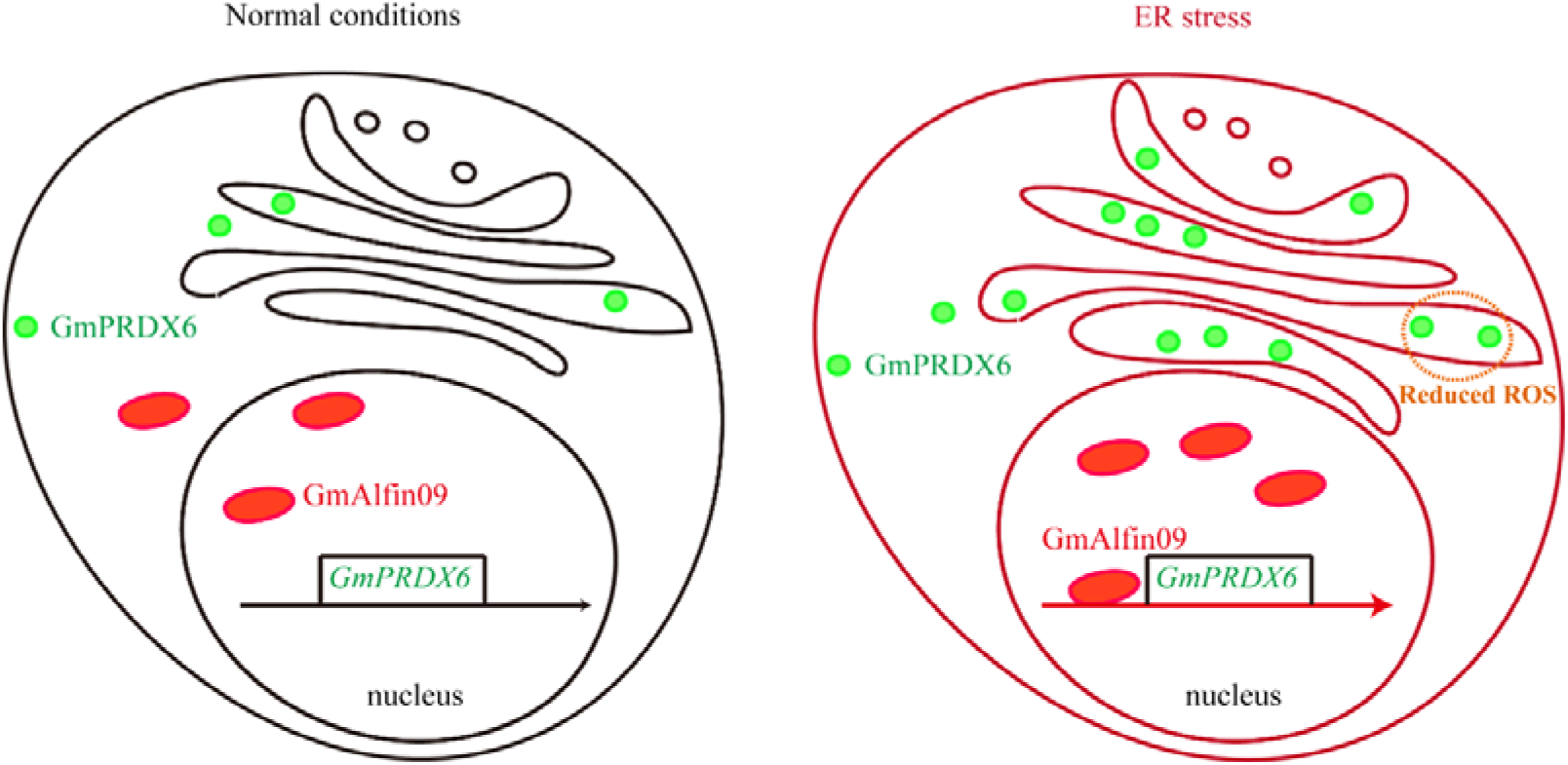
Model diagram of GmAlfin09-*GmPRDX6* module against ER stress. Under normal conditions, *GmAlfin09* and *GmPRDX6* expression is turned off. When ER stress occurs, GmAlfin09 enters the nucleus to start the expression of *GmPRDX6*, and the protein level of GmAlfin09 and GmPRDX6 increases. GmPRDX6 is largely enriched in the ER to clear the explosive electrons and ROS, and maintain cell stability.

Based on these results, we propose a hypothetical model for the function of this GmAlfin09-*GmPRDX6* regulatory module (Fig. 8). At baseline, only low levels of GmAlfin09 and GmPRDX6 expression are evident. In response to ER stress, however, GmAlfin09 localizes to the nucleus and promotes *GmPRDX6* expression, with corresponding increases in GmAlfin09 and GmPRDX6 protein levels. GmPRDX6 enrichment in the ER can then help clear the high levels of ROS and associated oxidative stress, helping cells remain viable and stable such that these soybean plants are better able to resist ER stress.

## DISCUSSION

### GmAlfin09 is a non-ER-anchored transcription factor that regulates ER stress responses

Most transcription factors to date that have been reported to be associated with ER stress are ER membrane-associated transcription factors. The plant AtbZIP17 and AtbZIP28 (Arabidopsis) and OsbZIP39 and OsbZIP60 (rice) proteins are ATF69 counterparts. In rice, the activated OsbZIP39, OsbZIP60, and OsbZIP50 proteins induce the expression of ER quality control (ERQC)-related proteins including the chaperone BiP9 (Hayashi et al., 2012). OPAQUE3 encodes a transmembrane bZIP transcription factor in rice that is involved in endosperm storage protein and starch biosynthesis (Cao et al., 2022), while the membrane-associated transcription factor bZIP28 detaches from the membrane and translocates to the nucleus to coordinate the unfolded protein response (UPR) (Sun et al., 2015). AtbZIP60 encodes a 295 amino acid protein with a predicted C-terminal transmembrane as well as a bZIP domain. Truncated AtbZIP60 isoforms lacking this transmembrane domain (AtbZIP60-C’) localize to the nucleus, consistent with the nuclear translocation of this protein after it dissociates from the membrane (Iwata and Koizumi, 2005). The AtNTL7 fragment can also be cleaved from the membrane of the ER in response to ER stress, localizing to the nucleus and thereby promoting the upregulation of target genes responsive to such ER stress (Chi et al., 2017). OsHLP1 can also coordinate with the ER membrane-associated NAC transcription factor OsNTL6 in rice cells to modulate ER homeostasis (Meng et al., 2022).

In the examples detailed above, transcription factors must be cleaved from the ER before they can enter the nuclear compartment. In contrast, GmAlfin09 was herein found to localize to the nucleus of soybean cells constitutively (Fig. 3C). Owing to the lack of any requirement for proteolytic cleavage, it may be able to respond to ER stress more directly, distinguishing GmAlfin09 from other membrane-related transcription factors.

### The GmAlfin09-GmPRDX6 axis serves as a different regulatory mechanism that alleviates ER stress-associated ROS generation

Alfin transcription factors can serve as positive regulators of stress resistance in plants. For example, when *Alfin1* is overexpressed in alfalfa, this can lead to the upregulation of the salt-induced *MsPRP2* in roots, consistent with the ability of Alfin1 to function as a regulator of root gene expression (Bastola et al., 1998). Overexpressing *Alfin1* in certain plants can thus enhance salt tolerance and growth. Most *BrAL* promoter regions contain cis-acting elements related to defense and stress responses (TC-rich repeats), and these genes were significantly upregulated in response to salt stress consistent with a role for these *BrALs* as regulators of salt stress responses in Chinese cabbage (Kayum et al., 2015). Transgenic *Arabidopsis* plants overexpressing the *GmPHD2* also exhibit superior tolerance for salt stress relative to wild-type plants (Wei et al., 2009).

Alfin-like transcription factors can also function as negative regulators of stress resistance in plants. For example, of the four *AhAL* genes identified in spinach, only *AhAL1* enhanced the salt and drought tolerance of *Arabidopsis* when constitutively expressed, whereas the other three increased *Arabidopsis* sensitivity to salt stress, highlighting clear functional differences among the members of this AhAL protein family (Tao et al., 2018). Similarly, adult-stage rice plants overexpressing *OsAL7.1* and *OsAL11* exhibited poorer drought tolerance (Yang et al., 2022). Seven total AL proteins have been reported in *Arabidopsis.* Despite their similar amino acid sequences, the stress responses of plants bearing mutant isoforms of these different proteins differ markedly. For example, the stress sensitivity of *al5* plants is substantially increased whereas *al6* plants are more stress-resistant. *AL7* overexpression can further inhibit the growth of *Arabidopsis* roots when these plants are exposed to salt stress such that *al7* mutant plants exhibit longer roots than those of control plants under salt stress conditions consistent with the role of *AL7* as a negative regulator of salt tolerance in these plants. *AL3* similarly serves as a negative regulator of plant salt tolerance, whereas *AL5* enhances this salt tolerance, further highlighting the distinct functional roles that these ALs play as regulators of stress resistance in plants (Tuller et al., 2013; Wei et al., 2015). In the present report, ER stress induced the upregulation of *GmAlfin09*. Overall, these findings suggest that Alfin genes can play dual roles in plant cells, with different members of this family exhibiting distinct functional roles that help coordinate plant stress responses. NbAL7 is a transcription inhibitor located in the nucleus, which can bind to the G-box motif in the promoter region of ROS scavenging genes such as *APX2*, *GR* and *GPX2* and inhibit their transcription, promoting ROS accumulation and enhancing resistance to TMV infection (Zhang et al., 2023).

However, in the present report, GmAlfin09 was found to act as a positive regulator and can directly promote *GmPRDX6* upregulation, with the resultant GmPRDX6 protein then being targeted to the ER where it functions as an ROS scavenger. This regulatory module thus represents a different mechanism through which soybean plants help alleviate ER stress and mitigate potential oxidative damage.

## SUPPLEMENTARY DATA

The following supplementary data are available at JXB online.

**Supplementary Figure S1.** Gene location, protein structure analysis and expression change.

**Supplementary Figure S2.** Prokaryotic expression and purification of GST-GmAlfin09 fusion proteins.

**Supplementary Figure S3.** Characterization of gene *GmPRDX6*.

**Supplementary Figure S4.** Analysis of promoter binding elements in *GmPRDX* gene family.

**Supplementary Figure S5.** The detection of transgenic lines in Figure 6A was performed.

**Supplementary Figure S6.** The positive detection of transgenic OE lines in Figure 7A was performed through PCR.

**Supplementary Table S1.** Primers used in the study.

**Supplementary Table S2.** Expression data of *GmAlfin* gene family.

## ACKNOWLEDGEMENTS

We would like to thank Professor Qingxin Song of the College of Agriculture of Nanjing Agricultural University for providing the genetic material of soybean mutant *gmalfin09*. We also want to thank MJEditor (www.mjeditor.com) for its linguistic assistance during the preparation of this manuscript.

## AUTHOR CONTRIBUTIONS

Kai Chen carried out experiments and wrote papers. Dongdong Guo, Jiji Yan, Huijuan Zhang, Zhang He, Chunxiao Wang and Wensi Tang participated in some experiments. Yongbin Zhou, Jun Chen and Zhaoshi Xu provided suggestions for writing the article. Ming Chen and Youzhi Ma provides guidance in experimental design.

## Abbreviations

transcription factor: (TF)
(ER stress): endoplasmic reticulum stress
(ROS): reactive oxygen species
(PRDX): peroxidase
(DTT): dithiothreitol
(EMSA): Electrophoretic Mobility Shift Assay

